# A Reproducible MFASS Benchmark of Splice-Disruption Predictors Reveals a Shared Exon-Interior Blind Spot

**DOI:** 10.64898/2026.07.21.739871

**Authors:** Brhanu F. Znabu, Zohaib Atif, Pradeep Devkota, Rediet Habtu Alemu, K.C. Jackfin

**Affiliations:** Biomedical Engineering Program, College of Engineering, University of Nebraska-Lincoln, Lincoln, NE, USA; Department of Biomedical Science and Engineering, Gwangju Institute of Science and Technology (GIST), Gwangju, Republic of Korea; Department of Molecular and Integrative Physiology, University of Kansas Medical Center, Kansas City, KS 66160, USA; Department of Biomedical Engineering, Kumoh National Institute of Technology, Gumi 39253, Republic of Korea; Department of Biological Systems Engineering, University of Nebraska-Lincoln, Lincoln, NE, USA

**Keywords:** RNA splicing, splice-disrupting variants, variant effect prediction, MFASS, multiplexed reporter assay, benchmarking, SpliceAI, Pangolin

## Abstract

We benchmark four published splicing variant-effect predictors against a multiplexed experimental splicing assay. On 27,733 single-nucleotide variants in and around human exons from MFASS with measured exon-inclusion outcomes, Pangolin is the strongest predictor of splice-disrupting variants (AUROC 0.888, average precision 0.421), ahead of SpliceAI (0.819, 0.321) and SpliceTransformer (0.786, 0.317), with MMSplice fourth (0.758, 0.256); all four exceed the older SPANR model (0.748, 0.228). The ranking reproduces the relative performance reported by the Pangolin authors, a correctness check on the pipeline. A calibrated consensus of the three deep-learning sequence-window predictors, evaluated on an exon-grouped held-out split, does not meaningfully improve over Pangolin alone. Stratifying by distance to the splice site exposes a shared blind spot: all five tools detect disruptions within a few bases of the splice site well, but recall declines sharply in the exon interior, and 19% of disrupting variants are missed by every tool; these shared misses are enriched among variants away from splice sites, and are predominantly exon-interior. MMSplice, the one model built for modular exonic and intronic effects rather than splice-site recognition, shows the same distance-dependent decline, so the blind spot is not an artifact of splice-site-centric architectures. Every number is computed against fixed experimental ground truth and is reproducible from the public dataset and the released code.

## 1 Introduction

RNA splicing is disrupted in a large share of disease-causing variants, and splice-altering variants are among the hardest to interpret in clinical genetics, because their effect depends on sequence context that is not obvious from the variant alone [1, 2, 3]. Deep-learning predictors of splicing from primary sequence, notably SpliceAI [4] and Pangolin [5], are now widely used to flag candidate splice-disrupting variants and appear in clinical variant-interpretation pipelines [6, 7].

These predictors, together with the more recent SpliceTransformer [8], the modular MMSplice model [10], and the earlier SPANR model [9], are usually evaluated in their own publications on heterogeneous datasets and metrics, which makes it hard to compare them on equal footing or to know where each is reliable. Multiplexed experimental assays of splicing provide a fixed ground truth for such comparisons [11]: MFASS (the Multiplexed Functional Assay of Splicing by Sort-seq) [12] measured the splicing effect of tens of thousands of human single-nucleotide variants in exons and their flanking intronic regions in a minigene reporter and labels those that strongly reduce exon inclusion as splice-disrupting variants.

Prior work has benchmarked deep-learning splice predictors against functional splicing assays [13, 14], and the MFASS variants themselves were used to compare splice scores when CADD-Splice was developed [15]. In one such effort, predictor concordance with the assay, and among predictors, was lower for exonic than for intronic variants [14]. Here we provide a uniform, reproducible, open benchmark of four modern predictors, together with the older SPANR model as a legacy baseline, on the MFASS single-nucleotide variants, scored through a single interface, with bootstrap confidence intervals and an exon-grouped held-out consensus. Beyond the aggregate ranking, we ask where the predictions fail: we stratify detection by distance to the splice site and identify the variants that every tool misses. Relative to prior MFASS-based comparisons [14, 15], the contribution here is twofold: a single-interface head-to-head that adds Pangolin, SpliceTransformer, and MMSplice with bootstrap confidence intervals and an exon-grouped held-out consensus, and a characterization of the shared exon-interior blind spot through the set of variants that every tool misses. We do not benchmark CADD-Splice itself, because it is a meta-score built on top of the sequence predictors evaluated here, so it would not be a like-for-like comparison.

## 2 Data and ground truth

MFASS measured the splicing effect of human single-nucleotide variants placed in exons and their flanking intronic regions in a minigene reporter [12]. Each variant carries a measured change in exon inclusion (the Δ inclusion index), and variants reducing inclusion by at least 0.50 are labeled splice-disrupting variants (SDVs). We use the public processed table, which provides hg38 coordinates, reference and alternate alleles, the continuous inclusion change, and the SDV label.

We score the 28,972 mutant single-nucleotide variants; 51% lie in exons and 49% in the flanking introns within about 50 bp of a splice site. For the binary detection task we drop variants whose v2 inclusion change was not measured (no label), leaving **27**,**733 variants, of which 1**,**050 (3.79%) are SDVs**. The strong class imbalance makes average precision (area under the precision-recall curve) the primary metric [17], reported alongside AUROC.

## 3 Predictors and scoring

We evaluate four published models using their released weights. Three are deep-learning models that score from a genomic sequence window: SpliceAI [4], a deep convolutional model; Pangolin [5], a tissue-aware successor; and SpliceTransformer [8], a transformer predicting tissue-specific splicing. Scoring for these three was orchestrated with the Proto tool ecosystem [16], but the models and weights are the published ones and the per-tool scores are released, so the benchmark is verifiable independently of that infrastructure. The fourth model, MMSplice [10], takes a different, modular approach: it scores the acceptor, exon, and donor sub-sequences of an exon separately and combines them, and was itself designed for and evaluated on modular exonic and intronic effects. We score MMSplice through its standard VCF interface against hg38, reconstructing each MFASS exon as an internal (cassette) exon so that both of its splice-site modules are defined; each variant’s effect is reduced to the magnitude of the predicted change in exon inclusion (|Δ logit Ψ|), matching the single-magnitude reduction used for the other tools. The older SPANR model [9], whose precomputed scores ship with MFASS, is included as a legacy baseline. Because the three sequence-window predictors require about ten kilobases of genomic context, each variant is scored in its real hg38 context (reference versus alternate), following each tool’s own usage, not in the short minigene fragment.

For each tool we reduce its output to a single disruption-magnitude delta score: SpliceAI’s maximum delta over its four acceptor/donor gain and loss scores; Pangolin’s larger of maximum splice gain and maximum splice loss magnitude; SpliceTransformer’s maximum change in acceptor or donor probability between the reference and alternate sequence at the variant locus, using the tissue-agnostic splice-type channels; the magnitude of MMSplice’s predicted change in exon-inclusion logit; and the magnitude of SPANR’s maximum-tissue Δ index. Tissue-aware tools are used in their tissue-agnostic setting, since MFASS is an in-vitro minigene assay.

## 4 Results

| predictor | $n$ | AUROC | AP | AP 95% CI |
| --- | --- | --- | --- | --- |
| Pangolin | 27,733 | **0.888** | **0.421** | [0.386, 0.454] |
| SpliceAI | 27,733 | 0.819 | 0.321 | [0.293, 0.353] |
| SpliceTransformer | 27,733 | 0.786 | 0.317 | [0.285, 0.352] |
| MMSplice | 27,733 | 0.758 | 0.256 | [0.226, 0.285] |
| SPANR (legacy) | 27,663 | 0.748 | 0.228 | [0.202, 0.260] |

Confidence intervals are 1,000-sample bootstraps of the average precision. SPANR covers slightly fewer variants because its precomputed table omits 70 of the evaluated variants, of which only 2 are SDVs, so the reduced coverage does not affect its ranking. The same ordering holds under both the ROC view (Figure 1) and the precision-recall view (Figure 2), which is the more informative of the two at this 3.79% positive base rate.

**Figure 1:**
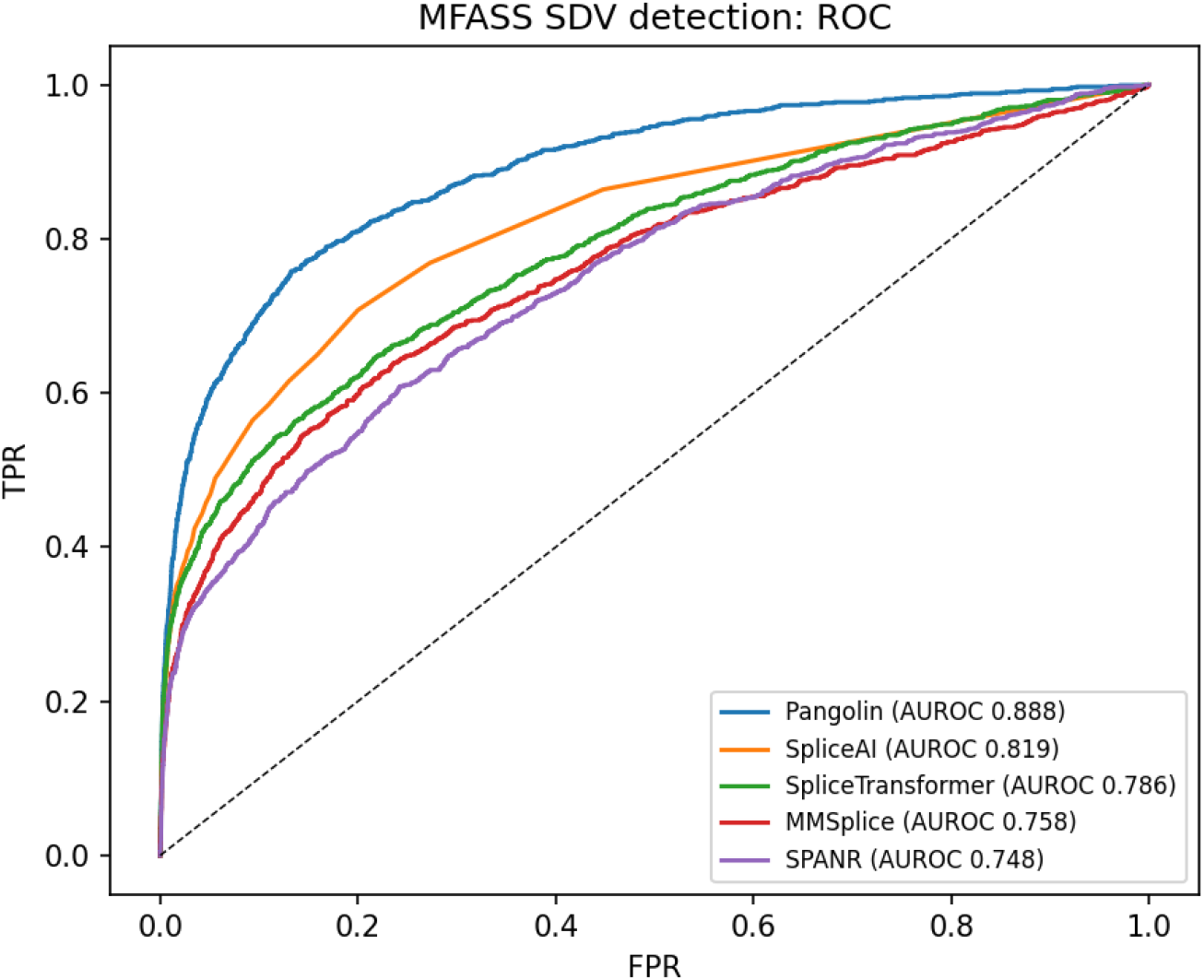
ROC for splice-disrupting-variant detection. Pangolin shows the strongest ROC performance; all modern tools exceed the legacy SPANR baseline.

**Figure 2:**
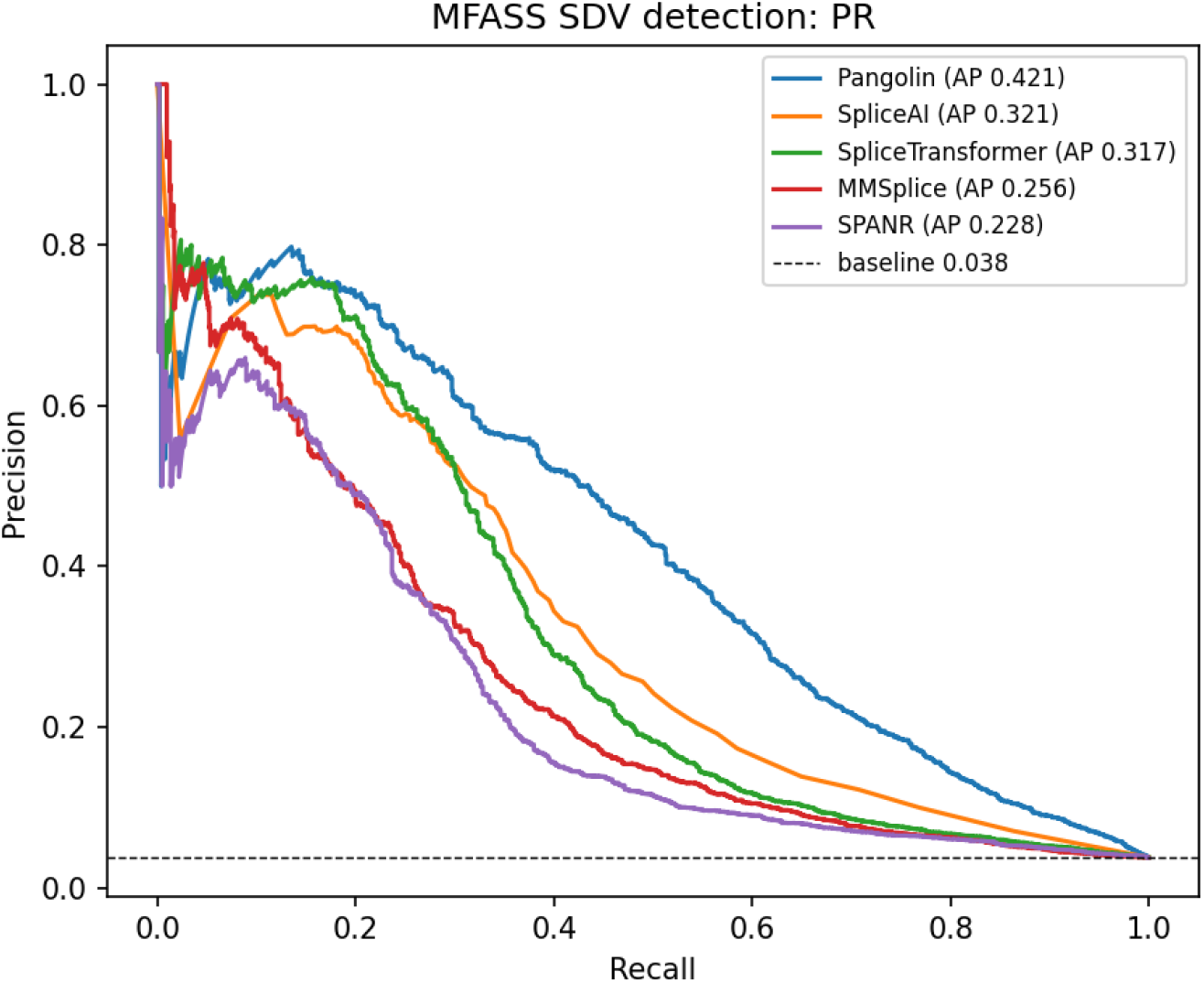
Precision-recall for the same task (dashed line is the 3.79% positive base rate). The ranking is consistent with the ROC view.

**Finding 1: Pangolin is the strongest predictor**, by a clear margin on both AUROC and average precision, and its AP confidence interval does not overlap those of the other tools.

**Finding 2: all four modern predictors beat the legacy SPANR model**, though MMSplice, used here tissue-agnostically on a single-cassette minigene context rather than a variant’s native multi-exon transcript, ranks fourth and sits just above the SPANR baseline.

**Finding 3: the ranking reproduces the relative ordering the Pangolin authors reported**, which is a correctness check on the scoring pipeline.

**Finding 4 (output granularity): in these outputs SpliceAI’s scores are more discretized than Pangolin’s**. SpliceAI reports its delta to two decimals and assigns exactly zero to most variants away from splice sites: 55% of non-disrupting variants but only 14% of SDVs score zero, so the zeros are genuine no-effect calls (zeros are depleted of SDVs, 0.96% versus the 3.79% base rate), not an artifact. Pangolin instead uses a continuous range, taking 27,528 distinct score values to SpliceAI’s 101, with no variant scored exactly zero. This finer score resolution may contribute to Pangolin’s stronger ranking here.

**Finding 5 (consensus): combining does not help**. A logistic-regression consensus of the three deep-learning sequence-window predictors, trained and evaluated on an *exon-grouped* 50/50 split (no exon appears in both train and test), reaches held-out AP 0.443 versus Pangolin’s 0.442 on the same split (both higher than the full-set 0.421 because this is the held-out half of the exon-grouped split). The difference of +0.0014 AP has a paired bootstrap 95% confidence interval of [*−*0.011, +0.012] that straddles zero, with 41% of resamples favoring Pangolin, and the consensus AUROC is slightly lower (0.890 versus 0.893). A rank-average consensus is worse (AP 0.372). The fitted model essentially relies on Pangolin alone (standardized coefficient 2.0, versus near zero or negative for the others). For this MFASS benchmark, Pangolin is the strongest single predictor.

## 5 Where the tools fail

The aggregate ranking hides where the predictions break down. We stratified splice-disrupting-variant recall by the distance of each variant to the nearest splice site, computed from the MFASS minigene construct annotation and cross-checked against an independent genomic-coordinate distance (Pearson *r* = 0.997), comparing all five tools at a common operating point: the score threshold that gives a 10% false-positive rate on non-disrupting variants.

Recall declines with distance from the splice site for every tool (Figure 3). Within 2 bp of a splice site the modern tools recover 80 to 90% of disrupting variants (Pangolin 0.90, SpliceAI 0.82, SpliceTransformer 0.80, MMSplice 0.85); beyond 20 bp their recall falls to 0.48, 0.35, 0.29, and 0.22 respectively, and SPANR to 0.20. More than half of the disrupting variants (53%) lie more than 10 bp from a splice site, and even Pangolin, the best tool, detects only 59% of them. This distance-dependent decline is independently consistent with the Pangolin authors’ own report, in which the model’s average precision falls from about 0.75 for variants within 9 bp of a splice site to below 0.35 beyond that distance [5]. MMSplice, despite its modular exon-centric design, follows the same profile as the splice-site models (high recall next to the splice site, falling into the exon interior) rather than resisting the decline.

**Figure 3:**
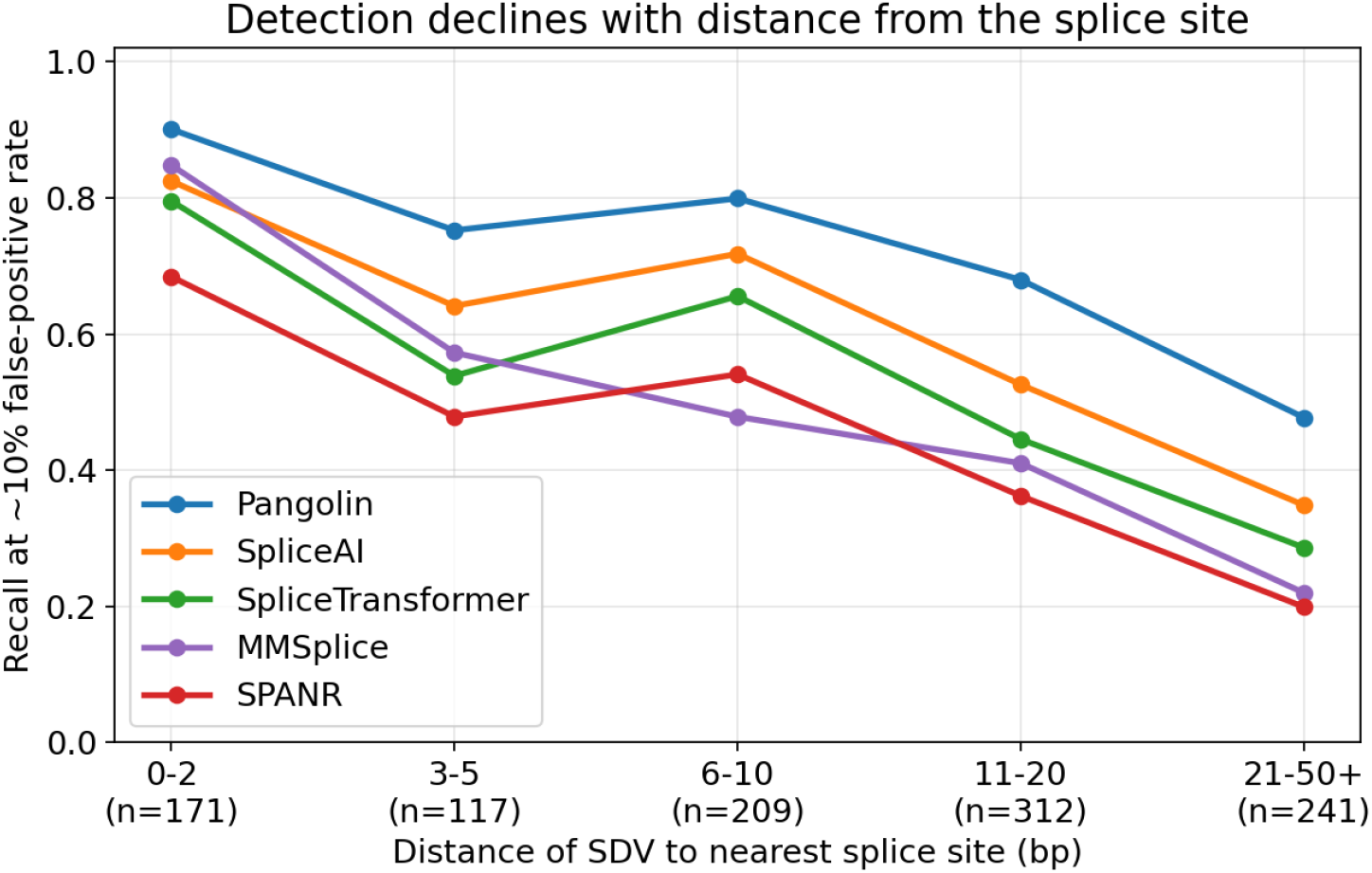
Recall of splice-disrupting variants, at a common 10% false-positive rate, as a function of distance to the nearest splice site. Every tool detects splice-site-proximal disruptions well and degrades with distance from the splice site.

Furthermore, 19% of all disrupting variants (200 of 1,050) are missed by every one of the five tools at this 10% false-positive operating point (30% at a stricter 5% and 8% at a more permissive 20% false-positive rate; the effect is qualitatively the same at every threshold). These shared misses sit predominantly away from the splice site: 77% lie more than 10 bp from a splice site, and 149 of 200 (74%) are exonic. They are not weak-effect variants sitting near the labeling cutoff: 82% have an inclusion change below *−*0.70, and only 4% sit within 0.1 of the *−*0.50 threshold. Their median change (*−*0.78) is modestly weaker than that of the detected disrupting variants (*−*0.87), but the shared miss is a true blind spot on strong-effect variants rather than an artifact of borderline labels. They are consistent with exon-interior regulatory disruptions, including possible exonic splicing enhancer or silencer effects [3], the class of variant MFASS was designed to expose. That MMSplice, which is explicitly built to model modular exonic and intronic regulation, misses these variants alongside the splice-site models indicates the blind spot is a shared empirical limitation of current predictors on this assay, not only a consequence of splice-site-centric design. The practical implication is that these tools are strongest for splice-site-proximal variants but require caution for exon-interior variants, where roughly one in five disrupting variants is invisible to all current predictors.

## 6 Scope and limitations

It shows that, against a multiplexed experimental splicing assay, Pangolin detects splice-disrupting variants better than SpliceAI, SpliceTransformer, MMSplice, and SPANR, that all four modern tools beat the legacy baseline, and that a simple consensus does not improve on the best single tool. Several caveats bound the claim. First, the predictors were trained on genomic and transcriptomic data and may have encountered the wild-type splice sites of these exons during training, so this is a realistic benchmark rather than a fully held-out one; this is standard for the field and applies to all tools. Second, MFASS is an in-vitro minigene assay in a single cellular context, so it does not capture tissue-specific or full-length-gene effects, and we therefore used the tissue-aware tools in a tissue-agnostic setting. Third, all predictors are scored as disruption magnitudes against a loss-of-function label; a directional or tissue-resolved analysis is left for future work. Fourth, MMSplice is scored on a reconstructed single-cassette minigene context rather than each variant’s native multi-exon transcript, and without a reference-inclusion term, which likely understates its absolute performance somewhat; its fourth-place ranking and its exon-interior decline should be read with that in mind. The contribution is a reproducible head-to-head benchmark, not a claim of clinical validation.

## 7 Reproducibility and data processing

The labels and continuous readout are the public MFASS measurements [12]; the three sequence-window predictor scores are produced by the public models run through Proto [16], and MMSplice by its public model through its standard VCF interface. Variants whose reported alleles are on the minus strand were normalized to the plus strand before scoring; for the three Proto-scored sequence-window predictors, full coverage of 28,972 of 28,972 variants confirms no variant was silently dropped. MMSplice likewise scores all 28,972 variants once each exon is presented as an internal cassette exon (an initial single-exon annotation left roughly half the variants, those in the flanking introns, unscored and is not used). SPANR covers slightly fewer because MFASS provides precomputed SPANR scores for only a subset. SpliceAI returns no score for variants outside an annotated gene; these were assigned a delta of zero, a minority as shown by the depletion of SDVs among zero-scored variants. This imputation can only depress SpliceAI, but it does not drive the ranking: restricting to the 12,855 variants SpliceAI scores nonzero (the three sequence-window tools and SPANR scored on the same subset), Pangolin still leads (AP 0.468) ahead of SpliceTransformer (0.369) and SpliceAI (0.365), with SPANR last (0.302). Every number is reproduced by `score_one_tool.py` and `score_mmsplice.py` (scoring) and `analyze.py` (metrics, consensus, and figures) from the public dataset.

## Supporting information

Supplemental Table 1: Benchmark metrics per predictor

Supplemental Table 2: Recall by distance to splice site

Supplemental Data: Full benchmark summary (JSON format)

## 8 Data and code availability

All code, the per-tool variant scores, the benchmark and failure-mode outputs, and the figures are openly available at https://github.com/brhanufen/spliceconsensus and archived on Zenodo (doi:10.5281/zenodo.20948820). The benchmark and all figures reproduce on a CPU in about a minute from the committed scores and a slim labels file, with no large download. The MFASS dataset is from Chong et al. [12] (https://github.com/KosuriLab/MFASS); the predictors are the public SpliceAI [4], Pangolin [5], SpliceTransformer [8], and SPANR [9] models, run through the Proto tool ecosystem [16].

## 9 Author contributions, competing interests, and funding

B.F.Z. conceived the study, designed the benchmark, implemented the analysis software, performed the analyses, generated the figures, and wrote the manuscript. Z.A. contributed to the study conception and revised the manuscript. P.D. contributed to validation of the results and revised the manuscript. R.H.A. contributed to the investigation and revised the manuscript. J.K.C. contributed to data curation and revised the manuscript. All authors read and approved the final manuscript. The authors declare no competing interests. No specific funding was received for this work.

## Notes

https://github.com/brhanufen/spliceconsensus

https://doi.org/10.5281/zenodo.20948820

### Competing Interest Statement

The authors have declared no competing interest.

